# Evidence of hatch-time based growth compensation in the early life history of two salmonid fishes

**DOI:** 10.1101/2022.06.06.495032

**Authors:** Heather D. Penney, Donald G. Keefe, Robert C. Perry, Craig F. Purchase

## Abstract

Initial body size can indicate quality within-species, with large size increasing the likelihood of survival. However, some populations or individuals may have body size disadvantages due to spatial/temporal differences in temperature, photoperiod, or food availability. Across-populations animals often have locally adapted physiology to compensate for poor environmental influences on development and growth, while within-populations behavioural adjustments that increase food intake after periods of deprivation provide opportunities to catch up (growth compensation). We posit a theoretical extension of growth compensation to include within-population differences related to short growing seasons due to delayed hatch time. We tested the hypothesis that individual fish that hatch later grow faster than those that hatch earlier. The relative magnitude of such a response was compared to growth variation among populations and between related species. We sampled young of the year Arctic charr and brook trout from five rivers in northern Labrador. Daily increments from otoliths were used to back-calculate size to a common age and calculate growth rates. Supporting the hypothesis, older fish were not larger at capture than younger fish, because animals that hatched later grew faster which may indicate age-based growth compensation.

## INTRODUCTION

Early phenotype can establish individuals on trajectories towards alternative life histories, and influences factors such as morphology, growth, and reproduction (Taborsky 2006; Jonsson and Jonsson 2014; Walsh et al. 2015; Rohde et al. 2015; Clarke et al. 2016). In turn, phenology (timing) of reproductive events such as germination, hatch, or birth affects early phenotypes (Beer & Anderson, 2001; Brännäs, 1995; Einum & Fleming, 2000; Sternecker, Denic, & Geist, 2014). Therefore, there is often strong selective pressure to reproduce at an optimal time (McNamara, Barta, Klaassen, & Bauer, 2011; Morgan & Christy, 1994; Morin, Lawler, & Johnson, 1990). For example, phenology has been shown to affect reproductive success in plants (Satake et al. 2001), corals (Guest et al. 2008; Mercier et al. 2011), insects (Maino et al. 2017), amphibians (Morin et al. 1990), fishes (Morbey and Ydenberg 2003), birds (Reed et al. 2009; Shoji et al. 2015), and mammals (Rotella et al. 2016). The fitness outcomes of variable timing of reproductive events are connected to subsequent growth conditions that offspring are likely to encounter.

Intra-specific variation in growth rate is ubiquitous. Among-population growth differences often exist due to latitudinal (and elevational) gradients in temperature and photoperiod, with individuals in higher latitudes experiencing shorter growing seasons (Campos et al. 2009; Sinnatamby et al. 2014). If phenotypic optima is similar across such conditions, disadvantaged populations can evolve greater genetic capacity for growth to mitigate some negative environmental effects on size via plasticity (Conover and Present 1990; Arendt and Wilson 1999; Purchase and Brown 2000; Campos et al. 2009; Pearson and Warner 2018). Such among-population patterns in local adaptation is termed counter-gradient variation (Figure 1).

**Figure 1.**
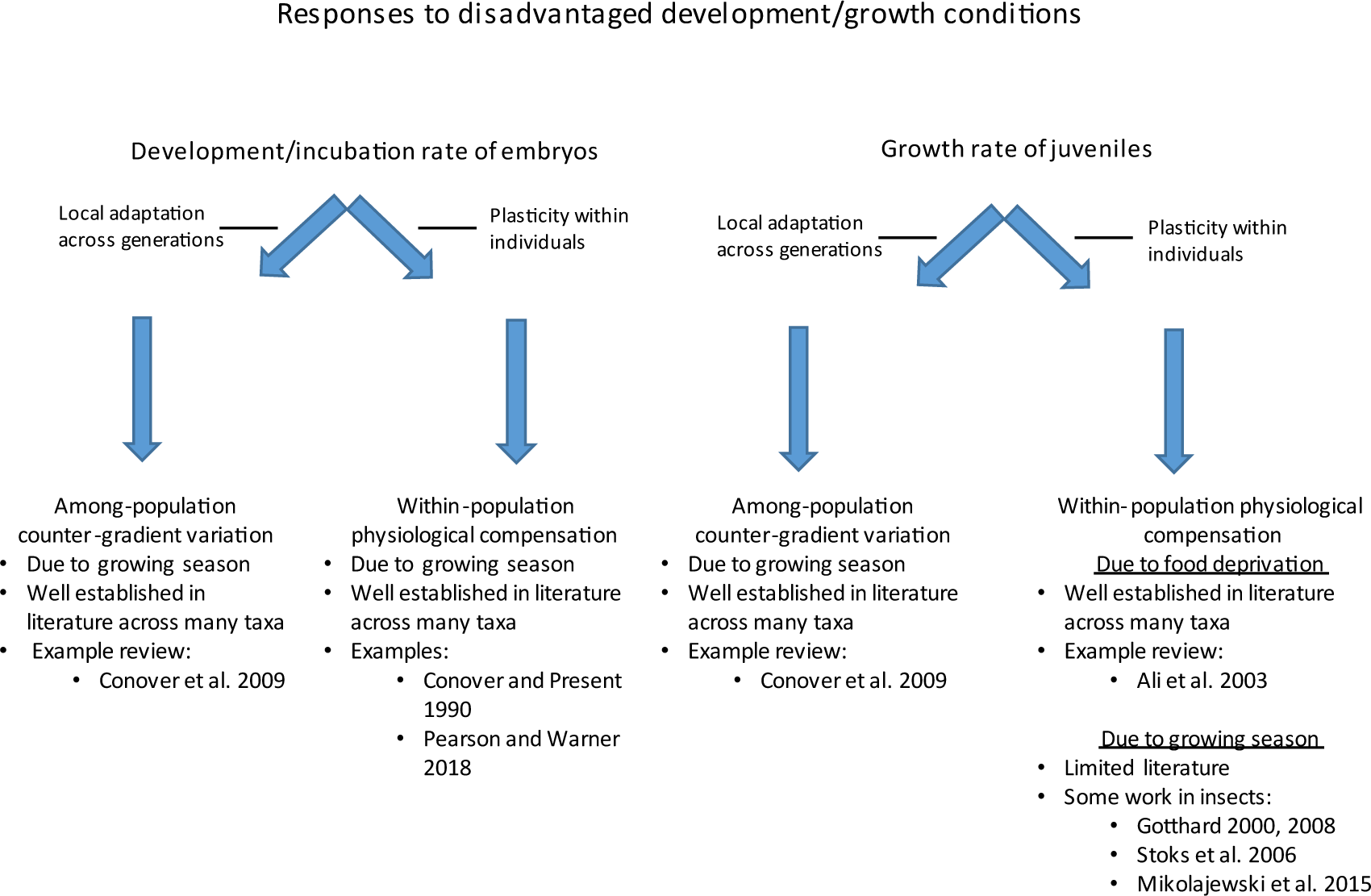
Schematic showing the theoretical development (embryos) and growth (juveniles) rate responses to disadvantaged conditions among-(local adaptation) and within-(physiological compensation) populations.

Within-populations, some individuals may experience relatively disadvantaged conditions. For example, a period of depressed feeding opportunities can result in diminished growth rates, however compensatory behaviours can allow them to catch up by the end of the growing season (Metcalfe and Monaghan 2001). This growth compensation (Figure 1) may be considered an intrinsic plastic response to changes in the environment (Zhu et al. 2003; Carlson, et al. 2004) that is triggered by environmental cues (Gotthard 2008), a depletion of energy stores (Ali et al. 2003), or timing of hatch (see below). Compensatory growth can have positive effects on individuals, through an increased likelihood of survival (associated with larger body size); however, growth compensation is associated with bolder foraging behaviours which put these individuals at a greater risk of predation (Nicieza and Metcalfe 1997; Damsgård and Dill 1998; Biro et al. 2004).

Among-populations, in addition to growth potential, counter-gradient variation in local adaptation has been associated with developmental timing (Figure 1); the concept is easily transferable. Both are influenced by temperature, and locally adapted physiology can produce common phenotypes despite substantial plasticity. Salmonid fishes provide a useful illustration. Salmonids are poikilothermic meaning their developmental rate is linked to temperature, hatching faster in warmer water. However, at colder temperatures they need fewer accumulated thermal units (ATU) to hatch (Brannon 1987; Quinn 2005), and there is evidence of counter-gradient variation of ATUs to hatch across populations (Sparks et al. 2019).

In contrast, within-populations, processes regulating developmental timing and subsequent growth rates have not been linked (Figure 1). Within-population hatch or birth timing is affected by environmental conditions (McNamara et al. 2011; Rooke et al. 2019), mating timing (Sternecker et al. 2014), parental genetics (Solberg et al. 2014), maternal condition (Berejikian et al. 2014) and investment in offspring (Beacham et al. 1985; Maino et al. 2017). Sub-optimal hatching phenology can result in a mismatch in trophic dynamics with prey (Brännäs 1995), whereby food is unavailable to newly hatched offspring. Optimal hatching timing is stochastic year to year but may be relatively stable across generations. Hatch too early and there may be no food and/or sub-optimal environmental conditions. However, late hatchers are at a competitive disadvantage for feeding territories compared to early hatchers due to dominance hierarchies (Metcalfe and Thorpe 1992; Cutts et al. 1999). Thus, sub-optimal hatch timing can result in slower growth rates and lower chances of survival (Snucins et al. 1992; Einum and Fleming 2000; Borcherding et al. 2010; Skoglund et al. 2012).

Previous work has established that growth rate is related to hatch time across-populations (e.g., Lapolla, 2001), and within-individuals, periods of slow growth due to limited food will be compensated by periods of faster growth when food availability increases (Metcalfe and Monaghan 2001). In this study, we posit a theoretical extension of growth compensation to include within-population differences related to shorter growing seasons due to delayed hatch time (Figure 1). An individual may compensate for hatching late, where they are disadvantaged by a shorter growing season, by growing faster than other individuals (within their population) that hatched earlier, thereby making the best of a bad situation. Such work has been supported by data from several insect studies (e.g. Gotthard 2000; Stoks et al. 2006; Mikolajewski et al. 2015), however it has not been explored in any vertebrate. We tested this hypothesis in two sympatric salmonid species (*Salvelinus spp.*) where the relative magnitude of such a response was compared to growth variation between the species and across populations (five rivers) in northern Labrador, Canada.

## MATERIALS AND METHODS

### Environmental information

Sampling occurred on secondary and tertiary streams of five river systems in northern Labrador, Canada: Hebron River, Kamanatsuk Brook, Fraser River, Anaktalik Brook, and Igluvigaluk Brook (Figure 2; Appendix A1). Temperature loggers (*n=*4, HOBO TidbiT v2, UTBI-001) were installed during the spawning season in October 2012 and removed during the June 2013 sampling period at two sampling sites, Fraser and Anaktalik rivers; and one additional river: Ikadlivik Brook (Appendix A1). Loggers were fastened to rebar and firmly placed in riverbeds (Appendix A2). Salmonids often spawn in groundwater seeps having relatively steady flows of water at stable temperatures. The temperature loggers placed in Fraser River were in a spawning aggregation where redds were observed, while the loggers in Anaktalik River and Ikadlivik Brook were placed in the main flow of the river. However, the temperature estimates from our loggers are likely underestimates (through winter) compared to those experienced in the redds because the loggers were in the water column and not in the gravel (where salmonids lay their eggs, and when in the presence of groundwater seeps, tend to have more stable temperatures).

**Figure 2.**
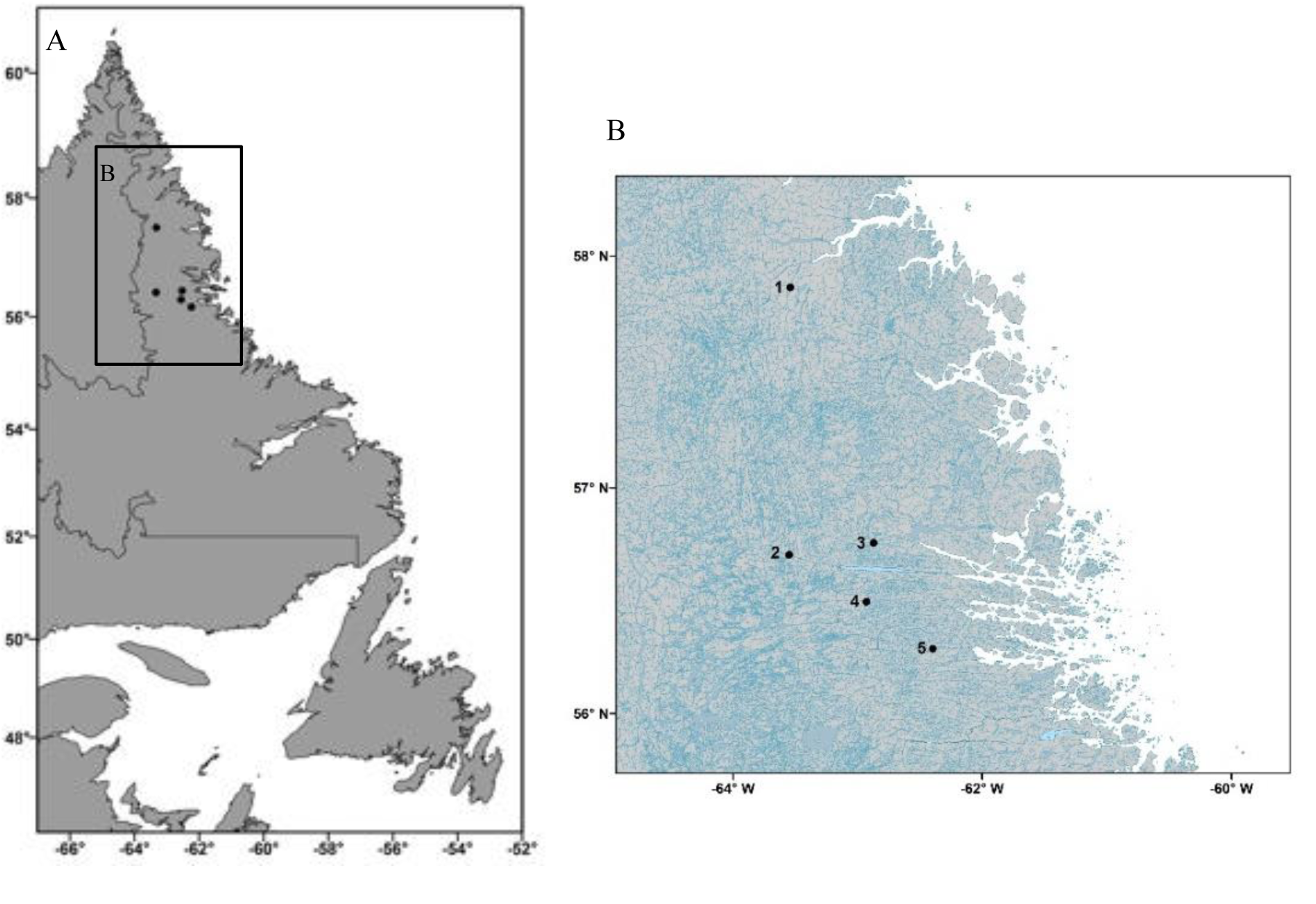
A) Map of Newfoundland and Labrador, Canada B) Inset of study area. For both maps, points indicate sampling sites, see Table A1 for site details and GPS locations.

### Fish collection

We collected young of the year Arctic charr and brook trout (*Salvelinus alpinus* and *S. fontinalis*) using a Smith-Root LR24 backpack electrofisher between June 24^th^ and June 29^th^, 2013. Potential collection sites were viewed from a helicopter with each tributary selectively sampled by electrofishing upstream. Areas where we were unlikely to find young of the year, such as sandy substrate, turbulent water, or water that was deeper than ∼1 meter, were not surveyed. The minimum stretch of water sampled was ∼100 meters per stream, and we only sampled two or three individuals from any one cluster of fish to minimize the risk of collecting multiple siblings from a family. After capture, the fish were euthanized, measured (fork length), and a tissue sample taken and placed in 95% ethanol for genetic species identification. The fish were then frozen (at -20°C) for later otolith extraction.

All animals were handled in accordance with the Guide to the Care and Use of Experimental Animals (Memorial University Animal care permit: 12-08-CP) and permitting from Fisheries and Oceans Canada.

### Genetic identification to species

The small physical size of the newly emerged hatchlings made morphological species identification difficult. Therefore, we used genetic barcoding to determine the species identity of each fish (brook trout or Arctic charr; n=436). We extracted DNA from tissue samples using a Qiagen DNeasy Blood and Tissue Kit according to the manufacturer’s protocol (Qiagen, Hilden, Germany). A 520 base pair fragment of the cytochrome oxidase 1 (CO1) gene was amplified by PCR using standard COI barcoding primers (Cox1-1F AACGTAATTGTCACCGCCCATG and Cox1-1R CACCTCAGGGTGTCCGAAG-AAT). We purified the PCR products with an Exo-SAP clean-up method and sent them to Genome Quebec (McGill University, QC) for sequencing using standard dideoxy methods. We aligned the sequences in MEGA v6.0 (Tamura et al. 2013) and species identification was unambiguously determined for all 436 fish.

### Otolith work and hatch date

Sagittal otoliths were extracted from young of the year fish using established methodology (Radtke 1996). Each otolith was fixed to a glass slide and polished using 3 and 30 μm lapping film. In salmonids a layer of calcium carbonate is deposited every day; this forms an increment that can be used to interpret fish age (see Radtke 1989, 1996; and Adams et al. 1992 for methods). When a band appears darker and thicker than the others, it is considered a check. Checks can occur for a variety of reasons including: stress due to hatch, emergence, or environmental change such as a storm, lack of food, or handling stress in aquaculture settings (Adams et al., 1992; Campana & Neilson, 1985). The result of this stress is slower growth, and therefore two or more daily rings merge into one thicker ring (Adams et al. 1992). In this study, we were interested in both hatch and emergence checks. The hatch check is a thick ring that encircles all of the primordia (nuclei upon which the otolith is built) near the core of the otolith. The emergence check occurs when hatchlings leave their gravel nest and begin exogenous feeding (Appendix A3). After the establishment of the emergence check, growth often accelerates and therefore subsequent rings are further apart and more translucent (Campana, 2001; Campana & Neilson, 1985). Other work has shown that hatch and emergence times can be inferred in wild salmonids based on otolith microstructure (Fitzgerald et al. 2021).

Of the original 436 individuals, only fish from rivers with a sample size n >10 young of the year fish of a species, and for which we could obtain daily age readings were included in further analyses (324 fish: 206 Arctic charr; 118 brook trout) (see Appendix A4 for details). We were unable to age 112 fish due to otolith loss or breakage during processing. Based on the number of daily increments present and the date of capture, each fish’s hatch date was back-calculated (Table 1). Photographs of otoliths were taken using a compound microscope under 100x magnification. The photographs were cropped, grey-scaled, and the colour range of greys was reduced to make ring visualization easier using Photoshop.

**Table 1.** Information for species in each site collected June 24 to June 29^th^, 2013 in Labrador, Canada. Site number refers to locations in Figure 1. Descriptive statistics for: mean age (in days, on June 24^th^ – first day of electrofishing), hatch date, mean otolith radius at capture (mm), fork length at capture, and estimated fork length at 25 and 50 days old (mm). Only samples from species in rivers that had a sample size of at least 10 individuals were included (N = sample size; SD = standard deviation).

| Species | Site | River | N | Mean ± SD* age (days) | Mean hatch date | Hatch Range | Otolith radius (mm) at capture ± SD* | Otolith radius (mm) at hatch ± SD* | Mean ± SD* capture fork length (mm) | Mean ± SD* estimated fork length at 50 days (mm) | Mean ± SD* estimated fork length at 25 days (mm) |
| --- | --- | --- | --- | --- | --- | --- | --- | --- | --- | --- | --- |
| Arctic charr | 1 | Hebron | 83 | 60±10 | Apr 28 | Mar 21-May 13 | 0.17±0.03 | 0.13±0.03 | 27.8±2.2 | 25.8±1.7 | 22.0±1.0 |
| | 2 | Fraser | 19 | 60±9 | Apr 27 | Apr 11-May 15 | 0.16±0.03 | 0.14±0.05 | 26.8±1.2 | 25.2±1.4 | 21.8±0.9 |
| | 4 | Anaktalik | 92 | 61±12 | Apr 24 | Mar 27-May 17 | 0.17±0.03 | 0.13±0.04 | 27.9±1.6 | 26.1±1.7 | 22.2±1.0 |
| | 5 | Igluvigaluk | 12 | 60 ±8 | Apr 20 | Apr 10-May 8 | 0.17±0.02 | 0.12±0.02 | 27.6±3.0 | 25.9±2.4 | 21.9±1.2 |
| Brook trout | 3 | Kamanatsuk | 100 | 58±8 | Apr 23 | Mar 30-May 17 | 0.17±0.03 | 0.13±0.04 | 26.8±2.7 | 25.2±3.2 | 21.7±1.8 |
| | 5 | Igluvigaluk | 18 | 59±11 | Apr 25 | Apr 9-May 8 | 0.17±0.02 | 0.13±0.03 | 25.0±3.8 | 23.8±2.3 | 21.0±1.1 |

\*SD= standard deviation

Each otolith was assigned a blind code and read without knowledge of species or river origin. We reinterpreted age on a random subset of 50 fish to determine precision. Precision estimates were based on coefficient of variation (CV) values (Chang 1982). A review found that a CV of less than ∼8% is generally acceptable for aging studies (Campana, 2001). Our precision estimate was 7.1% for days post-hatch and 8.0% for days post-emergence. However, when we compared the 40 emergence checks in the 50 subsampled otoliths to the original interpretation, 17.5% (*n=* 7) did not agree (i.e. a check was noted in one reading but not the other). Therefore, determining emergence checks was deemed unreliable for these otoliths, and we did not examine emergence time in further analyses. All otoliths were aged by the same reader (HDP).

### Growth rate and back calculated lengths

We used the ObjectJ plugin for ImageJ (Schneider et al. 2012) to calculate the fish’s daily growth rate (mm/day) based on the width of the daily otolith rings. The total radius of the otolith was measured from the hatch line to the last visible increment. Daily growth was then calculated based upon total growth of the fish (length at capture minus estimated hatch size; defined below) compared to the width of each daily otolith ring. We back-calculated hatchling length to 25 and 50 days post-hatch (Table 1) using the biological intercept model [model 1] (Campana 1990; Vigliola and Meekan 2009).

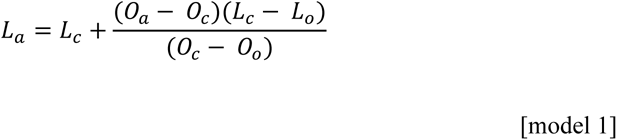

Where L was fish length (size, mm), O was otolith radius (µm), L_c_ and O_c_ were size at capture, L_a_ and O_a_ were size at age, and L_o_ and O_o_ were size at hatch. One of the weaknesses of this model was that hatch length needs to be estimated in order to estimate post-hatch lengths. We used a hatch length (L_o_) of 18 mm based on previous work in brook trout (Penney, Beirao, and Purchase, 2018). Additionally, we conducted a sensitivity analysis with an assumed hatch size of 16 mm and 20 mm and it made no difference on the overall conclusions.

We calculated each fish’s growth (mm) each day based on the width of each otolith ring. Next an average daily growth rate (average mm/day) was calculated based upon total growth of the fish (length at capture minus estimated hatch size, see model 1) compared to the width of each daily otolith ring. We compared growth rate (mm/day) for 3 different time periods. First, we examined the first 25 days of life to compare age effects on growth, next we examined a set time period (from June 1 to 21) to compare growth rates during standardized environmental conditions (photoperiod, and temperature), and finally we looked at overall growth rates. For each of these instances, we used model 1 to estimate growth rate.

### Data analyses and statistics

For descriptive purposes and to determine if we could examine our hypothesis at a species level, we tested whether river (population) had an effect on any of our dependent variables (hatch date, growth rate, fork length) (Table 2). To determine if hatch date (HD, taken from age on June 24^th^) or overall growth rate (GR, mm/day) differed between species (Sp) or among rivers (R) we conducted two analyses of variance (ANOVA) using model 2. Since there were no differences in growth rates in the first 25 days, the 3-weeks in June, or the overall growth rate we chose the overall growth rates in our analysis.

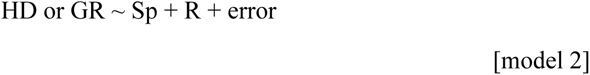

**Table 2.** Results of analysis of variance (ANOVA) for hatch date and average growth rate (mm/day) between fish species (*Salvelinus alpinus* and *S. fontinalis*) and analysis of deviance (**χ^2^)** for fork length (mm) for samples collected among five rivers June 24 to June 29^th^, 2013 in Labrador, Canada (df = degrees of freedom; F = calculated F statistic; p – probability; **χ^2^ =** analysis of deviance).

|  | Hatch date |  |  | Fork length (mm) |  |  | Growth rate (mm/day) |  |  |
| --- | --- | --- | --- | --- | --- | --- | --- | --- | --- |
| Factor | **F** | **df** | **p** | $\chi^2$ | **df** | **p** | **F** | **df** | **p** |
| Species | 0.21 | 1,318 | 0.65 | 4.98 | 1,318 | **0.03** | 4.21 | 1,318 | **0.04** |
| River | 0.16 | 1,318 | 0.96 | 7.38 | 4,318 | 0.12 | 1.99 | 4,318 | 0.10 |
| Age | NA | NA | NA | 3271.76 | 1,318 | **0.0001** | NA | NA | NA |

In addition, we ran an analysis of deviance on a linear mixed effect model (LME) to examine differences in fork length (mm) between species and among rivers at two points (25 and 50 days post-hatch) [model 3]. Where FL was fork length, A was age, Sp was species, and R was river. ID was a unique identifier that was a random factor which allowed for paired results between the two ages within an individual.

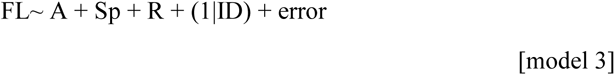

We did not test for the interaction between species and river for model 2 or 3 because we did not have representatives from both species in each river. There was no effect of river on hatch date (above), fork length at capture, or growth rate (Table 1, Table 2), therefore the populations were pooled for further analyses. Therefore, we conducted linear regressions for each species (regardless of river) to determine: 1) if there was an association between fork length at capture and age, and 2) if there was an association between hatch date and growth rate. For all analyses, α was set at 0.05. Residuals were examined to test for normality and heteroscedascity, and no deviations were observed. The map was created in ArcGIS. All graphs (ggplot2), data processing and statistics were done in R version 3.3.3 (R Development Core Team, 2015; using packages car, ggpmisc, Hmisc, lme4, and lubridate).

## RESULTS

The hatch dates (Table 1) for brook trout (mean: April 26; range: March 30 to May 17) and Arctic charr (mean: April 24; range: March 21 to May 17) did not differ between species, or among rivers (Table 2). Brook trout (26.5 ± 3.7 SD) were slightly shorter than Arctic charr (27.7 ± 1.8 SD), (Table 2, Figure 3). There was a significant difference in individual growth rates by species, with Arctic char growing faster than brook trout (Table 2).

**Figure 3.**
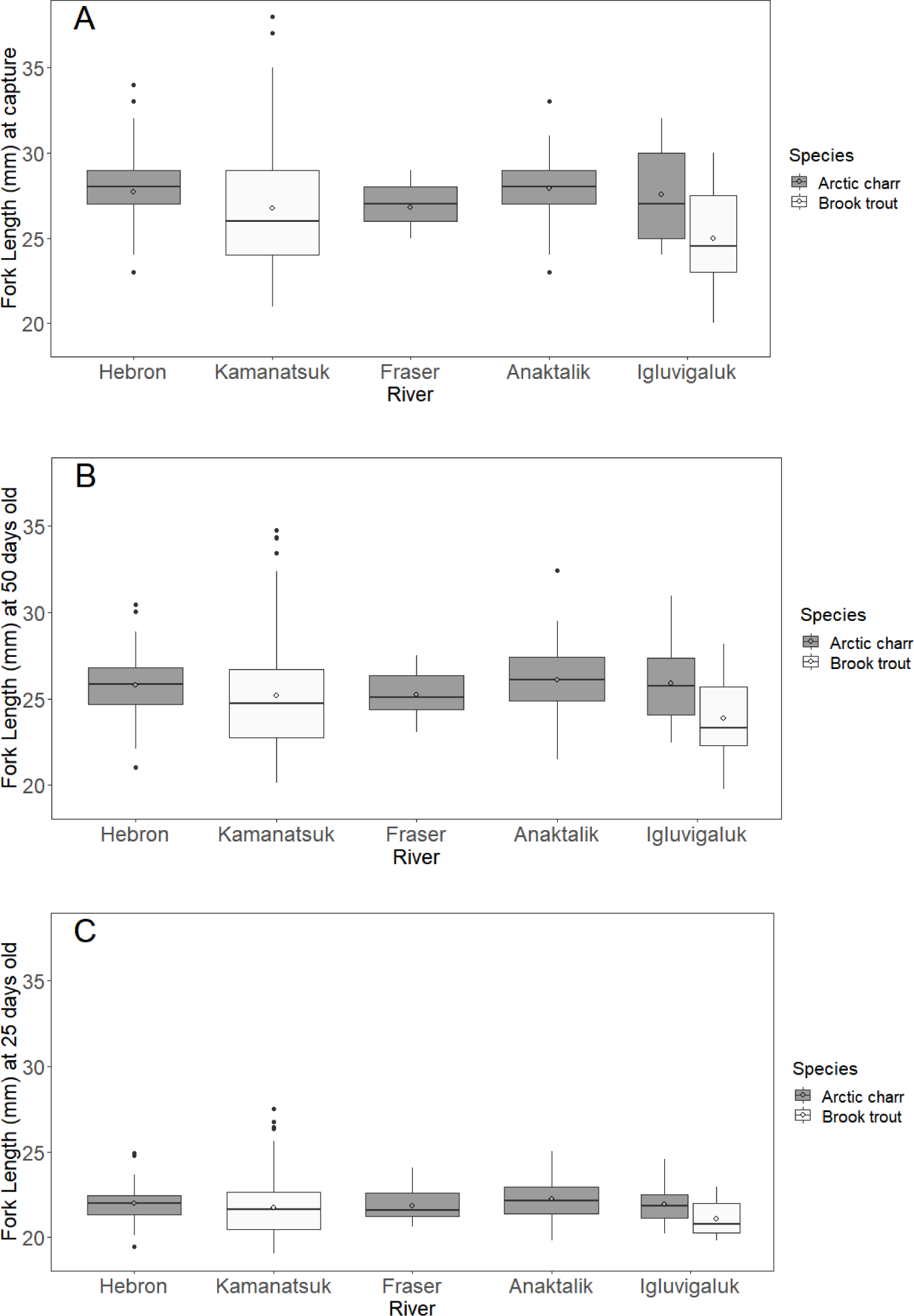
Box plot comparisons of fork length at (A) capture, and back-calculated fork lengths for (B) 50 and (C) 25 days old for Arctic charr and brook trout (*Salvelinus alpinus* and *S. fontinalis*) and river (only rivers with a sample size greater than 10 were included) sampled June 24 to June 29^th^, 2013 in northern Labrador, Canada. The boxplot shows the median (line) the interquartile range (IQR, 25 and 75%), whiskers represent the next quartile of the data (1.5 *IQR), and outliers are represented by dots. Open circles represent the mean for each group.

Subsequent to pooling populations, we conducted linear regressions and found no relationship between age and fork length at capture for Arctic charr or brook trout (Figure 4). Older fish were not bigger than younger fish. To better understand the absence of a relationship between age and body size at the same capture time, we conducted additional tests regressing hatch date to daily growth rate at three time points. We found significant relationships with all three time points (first 25 days, June 1^st^ to 21^st^, and entire post-hatch life) for both Arctic charr and brook trout (Figure 5), whereby, fish with earlier hatch dates had slower growth rates than fish that had later hatch dates.

**Figure 4.**
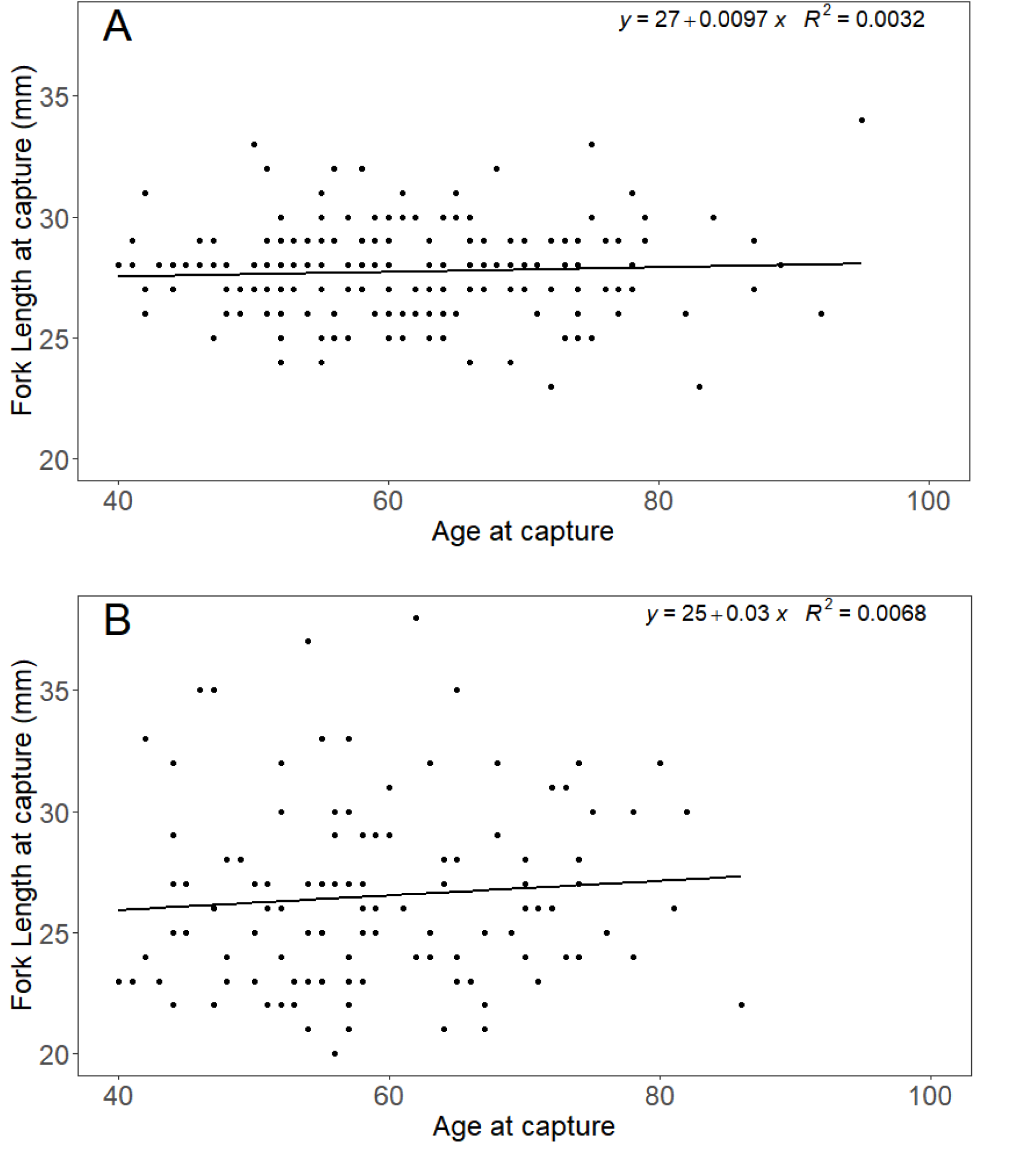
Plots showing lack of a relationship between age (days) and fork length (mm) at capture for (A) Arctic charr and (B) brook trout among five rivers in northern Labrador, Canada.

**Figure 5.**
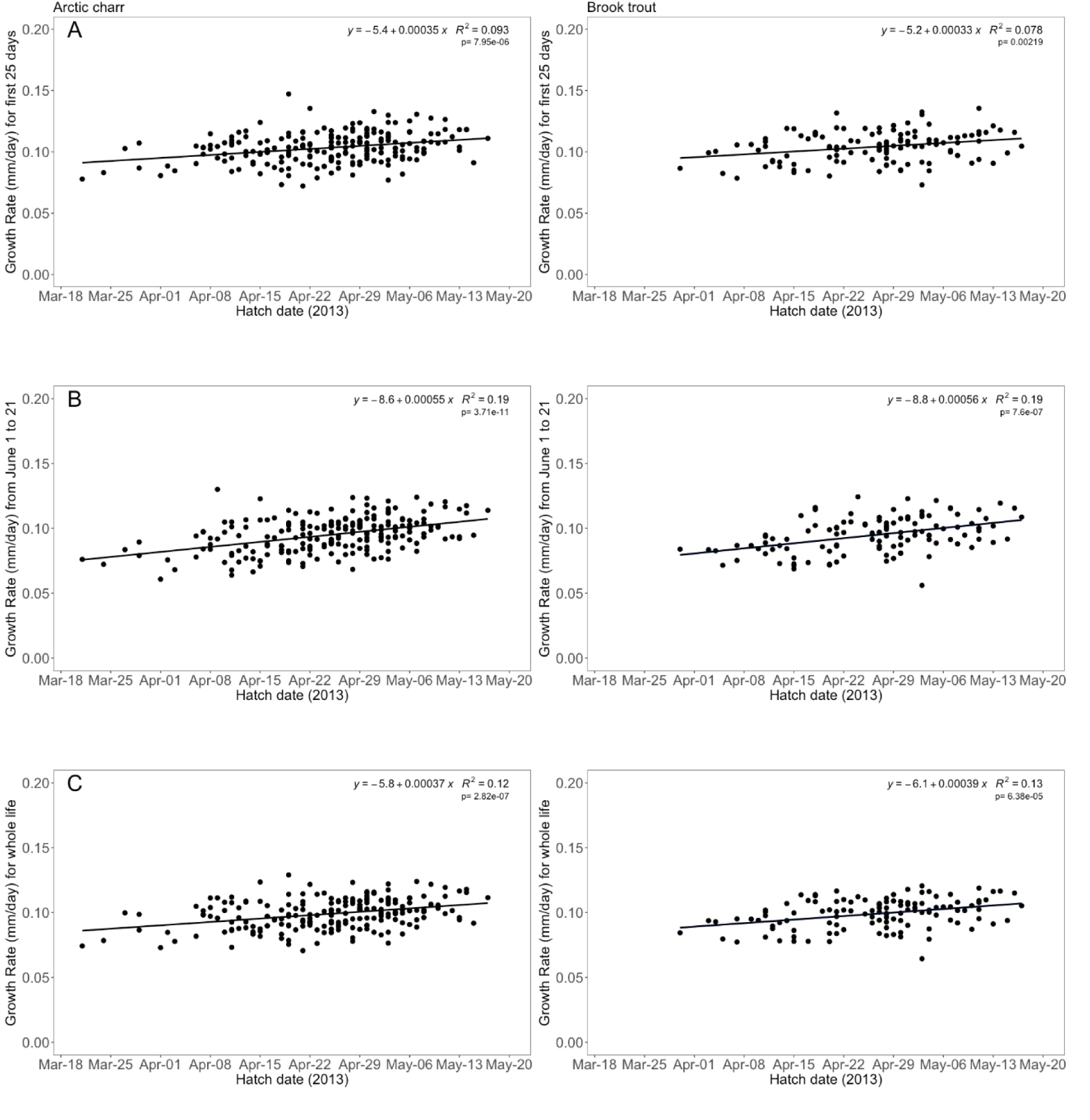
Plots showing simple linear regression models for the relationship between hatch date and average daily growth rate regardless of river for (A) the first 25 days of life; (B) the time period of June 1st to 21st; and (C) for their entire life post-hatching (average mm/day, based otolith estimation) for Arctic charr (left) and brook trout (right) (*Salvelinus alpinus* and *S. fontinalis*) among five rivers in northern Labrador, Canada.

## DISCUSSION

The objective of this paper was to test the hypothesis that hatch time affects growth rate, where individuals that hatch later are disadvantaged by a shorter growing season than those that hatch earlier and therefore would grow faster to compensate, potentially making the best of a bad situation. We found support for our hypothesis in two sympatric species of salmonids. Overall, there was more variation in growth rate among individuals within-populations, than across-populations or between species. There was no relationship between age (in weeks) and size of young of the year charrs (>75 days old, sampled in late June), which means that older hatchlings were not larger than younger ones. We found that this occurred because fish that hatched later grew faster, potentially as a form of growth compensation. Growth compensation has been found in other studies and has been linked to initially poor environmental conditions such as drought (Oesterheld & McNaughton, 1991), high density (Sundström et al. 2013), or low food availability (Metcalfe and Monaghan 2001; Walling et al. 2007). Thus, differences in growth rates due to hatch timing may be a theoretical extension of growth compensation within-populations as an adaptive response to a short growing season experienced by late hatchers, as shown in insects (see: Gotthard 2000; Stoks et al. 2006; Mikolajewski et al. 2015), however, to our knowledge has not been shown in vertebrates.

Within species, variation in growth rate occurs on population (counter-gradient variation; e.g., Carlson, et al., 2004; McCairns, 2004; Yamahira & Conover, 2002) and individual levels (growth compensation; e.g., Mortensen & Damsgård, 1993; Nicieza & Metcalfe, 1997). There is predictable latitudinal variation in temperature and photoperiod that contribute to counter-gradient variation in growth potential among populations, where higher latitude populations evolve faster growth rates at equivalent temperatures to that of those at lower latitudes (Lapolla 2001; Campos et al. 2009; Sinnatamby et al. 2014).

Individuals may have poor endogenous resource stores if they have few opportunities to gather resources. This situation can arise through a food shortage brought on by a competitive disadvantage from not establishing feeding territories before others in their cohort (Metcalfe and Thorpe 1992; Cutts et al. 1999), or shorter growing seasons (Arendt and Wilson 1999; Campos et al. 2009). A period of faster (compensatory) growth often occurs after a depletion of resources causes a period of slow growth (Metcalfe and Monaghan 2001; Ali et al. 2003). Individuals counteract or compensate for this disadvantage by growing faster, which allows them to reach a similar size to individuals who were not stunted at a later time point. To investigate this process in response to hatch timing instead of food, we examined growth rates at three time points: during initial growth, for the entire life post-hatch, and for the first three weeks of June. We found that during all three time periods individuals that hatched later grew faster than early hatchers. This phenomenon was observed in daily otolith growth increments, where incremental growth of late hatching individuals tended to be larger than that of early hatching individuals. The June growth comparisons indicate that late hatchers grow faster even under the same abiotic conditions and food availability (discussed below).

Previous work in other salmonid species has shown that larger eggs tend to produce bigger offspring, and larger offspring may emerge from the nest (i.e., ready to begin exogenous feeding) earlier than smaller offspring (e.g., Solberg *et al*., 2014; Cogliati *et al*., 2018). The probable difference in resource availability (both diminished fat stores and yolk resources) in early life may be enough to trigger a growth compensation response in the late hatching fish. Additionally, individuals that are larger have a higher absolute growth rate but a lower relative or proportional growth rate (Van Buskirk et al. 2017). Cogliati et al. (2018) found that when comparing early and late hatchers there was no difference in growth rate, but did find that fish from small eggs had a significantly larger increase in size over time. In our case, the older fish grew slower, therefore the potential bias is in a conservative direction because the small fish had a higher absolute growth rate and a higher proportional growth rate. We do not know whether intrinsic effects such as hatch time or extrinsic environmental conditions experienced at different hatch times were the main causes for differences in growth rate. There are likely notable differences in food supply (e.g., insect abundance increases through the spring) and photoperiod in the experiences of the early and late hatchers, which might explain the differences in growth rates. One could also assume that temperature would be an extrinsic explanation for the pattern of increased growth rate later in the season. However, when comparing the first 25 days of life experienced by early hatchers (early April, ∼1°C) and late hatchers (mid-May, ∼2°C), the temperature profiles were likely not different enough to explain the difference in growth rates. Most importantly, the comparison during the first 3 weeks of June controls for differences in both abiotic and biotic effects, and showed that the pattern was the same. Due to prior residence advantage for early hatchers (O’Connor et al. 2000) we speculate that in order for late hatchers to grow faster, they possibly had to feed longer and/or more aggressively which is contrary to previous work that generally shown that early emergers tend to be bolder (Laakkonen and Hirvonen 2007; Vaz-Serrano et al. 2011).

Our estimated hatch timing fits with estimated temperatures (∼1 to 2°C), because Labrador populations spawn in mid October to early November (DFO 2001) and we would expect a hatch range of late March to mid-May (∼160-190 days at 1-2°C). Previous work has shown that Arctic char hatch between 331 to 416 accumulated thermal units (ATU) (at 8.5°C, 39 to 49 days; Yanik, Hisar, & Bölükbasi, 2002) and brook trout between 477 to 483 ATU (at ∼10-11°C, 43 to 48 days; Witzel and MacCrimmon 1983, Penney et al. 2018). We found relatively similar results in lab experiments conducted on Arctic charr embryos from the Fraser River, Labrador (400±34.9 ATU; Penney and Purchase unpublished data).

The results of this study show that the timing of hatch affects growth rate, providing more evidence that hatch phenology can play an important role in early life history. Growth rate and size are two important early life history traits in fishes and understanding the nuances of factors that affect early growth can help explain how different early phenotypes project into different adult life history strategies (Clarke et al. 2016). Future work designed to empirically test specifically how hatch phenology affects growth rate and survival through the first year of life, and how that translates into differences in fitness for salmonids is recommended. Furthermore, predicted changes in climate are likely to affect hatch phenology (Rooke et al. 2019), and therefore, more research should be done to understand the consequences of changes in complex northern ecosystems.

## Acknowledgements

The authors would like to thank I.A. Fleming and I.R. Bradbury for their contributions in the early conception of the project, L. Lait for help with genetics, B. Devine for creating our map, S.E. Campana for training H.D.P. to read otoliths, and J.B. Dempson, I.A Fleming, C. Brown, M. Abrahams, S. Einum, L. Lait, and T. Van Leeuwen for comments on an earlier version of the manuscript. Funding was provided through grants to C.F.P. from the Forestry and Wildlife Research Division and the Research and Development Corporation of the Government of Newfoundland and Labrador, the Natural Sciences and Engineering Research Council of Canada, and the Canada Foundation for Innovation. H.D.P. was also supported by a scholarship from the Natural Sciences and Engineering Research Council of Canada.

## Appendices

**Appendix A1:** Fish sampling locations (degrees decimal minutes) and temperature logger locations sampled June 24 to June 29^th^, 2013, in northern Labrador, Canada. Number of loggers at each location included in brackets. (TL = locations with temperature loggers).

| Site | River name | Sampled | Coordinates |  | Logger location |
| --- | --- | --- | --- | --- | --- |
|  |  |  | Latitude | Longitude |  |
| 1 | Hebron River | Yes | 57°N 51.96 | 63°W 32.37 |  |
| 3 | Kamanatsuk Brook | Yes | 56°N 45.48 | 62°W 52.31 |  |
| 2 | Fraser River | Yes | 56°N 42.34 | 63°W 32.90 (TL) | Spawning bed (2) |
| 4 | Anaktalik Brook | Yes | 56°N 29.94 | 62°W 55.68 (TL) | River (1) |
|  |  |  | 52°N 30.03 | 62°W 56.79 |  |
|  |  |  | 56°N 30.01 | 62°W 55.37 |  |
|  | Ikadlivik Brook | No | 56°N 24.00 | 62°W 31.55 (TL) | River (1) |
| 5 | Igluvigaluk Brook | Yes | 56°N 17.43 | 62°W 23.86 |  |
|  |  |  | 56°N 17.64 | 62°W 23.58 |  |
|  |  |  | 56°N 16.64 | 62°W 26.06 |  |

**Appendix A2.**
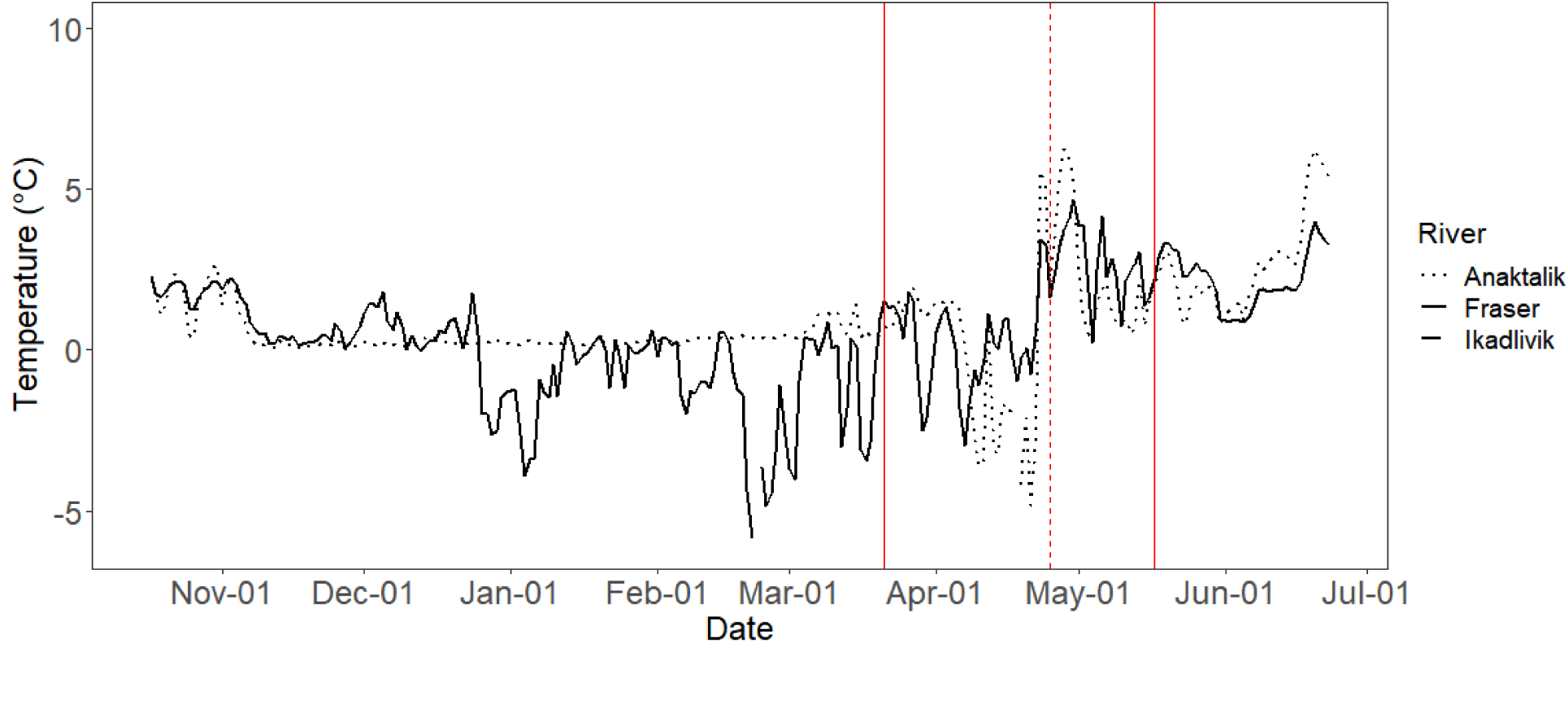
Temperature profiles from Fraser (black line) (average of 2 loggers), Ikadlivik (grey, dashed line) and Anaktalik (black, dotted line) rivers in Labrador, Canada from loggers in place from October 2012 to June 2013. Vertical lines indicate hatching dates for both species and all rivers (solid=range, dashed= mean). Note: negative values likely indicate being frozen in ice.

**Appendix A3.**
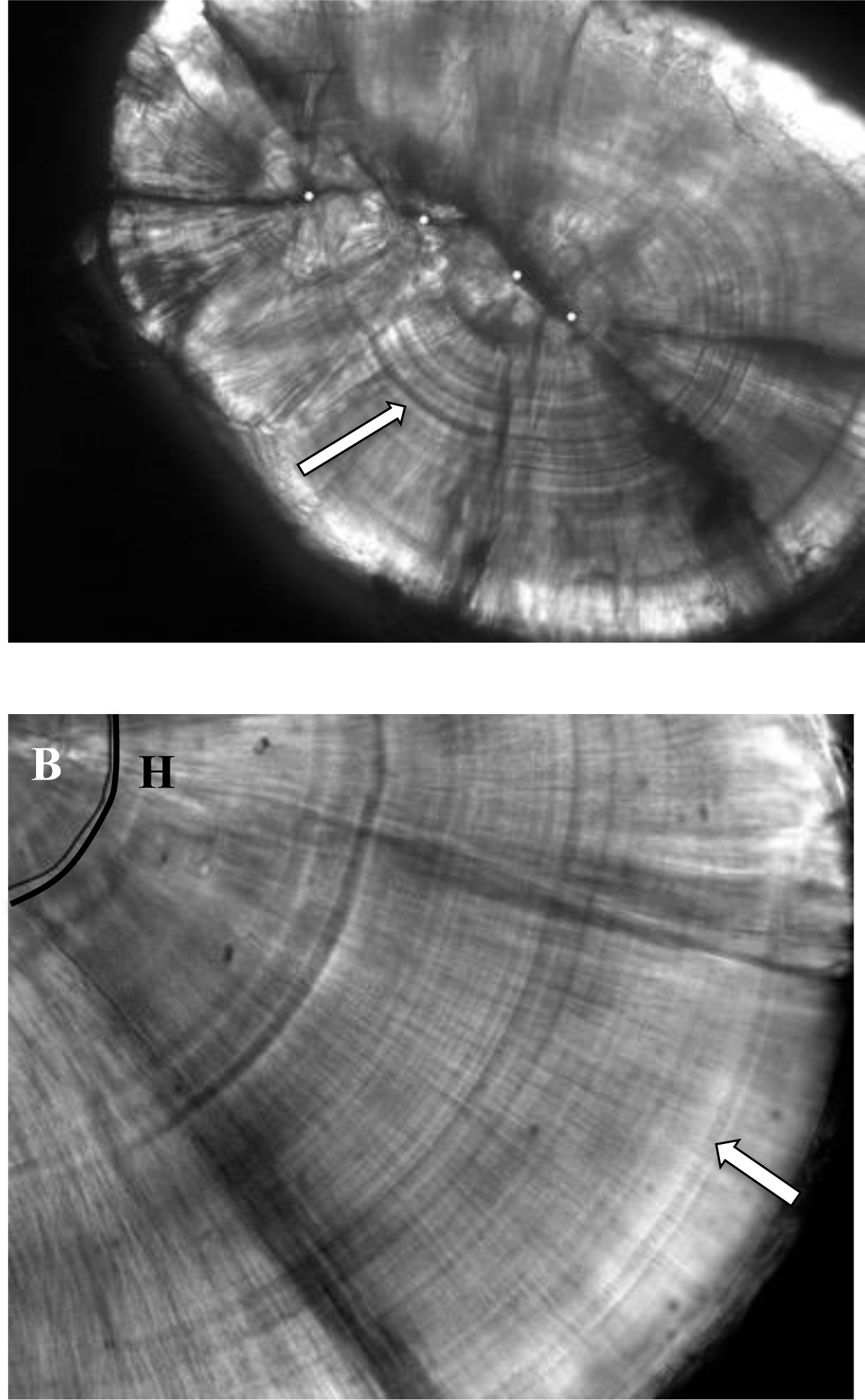
Photographs of otoliths from Arctic charr sampled from Anaktalik river, Labrador taken under a compound microscope and then manipulated in Photoshop. A) A whole salmonid otolith (40x), and B) a close-up photo (100x, under oil immersion), with hatch (H) and emergence (white arrow) checks indicated. White dots indicate primordia.

**Appendix A4:** Total number of Arctic charr and brook trout (*Salvelinus alpinus* and *S. fontinalis*) for each river caught June 24 to June 29^th^, 2013 in Northern Labrador, Canada and subsequently aged. Only species that had data on more than 10 individuals in a river were included in the analyses.

|  |  | Anaktalik | Fraser | Hebron | Kamanatsuk | Igluvigaluk | Total |
| --- | --- | --- | --- | --- | --- | --- | --- |
| **Arctic charr** | Caught | 100 | 40 | 107 | 3 | 18 | 268 |
|  | Aged | 92 | 19 | 83 | 0 | 12 | 206 |
| **Brook trout** | Caught | 5 | 3 | 4 | 129 | 27 | 168 |
|  | Aged | 0 | 0 | 0 | 100 | 18 | 118 |
| **Total** | Caught | 105 | 43 | 111 | 132 | 45 | 436 |
|  | Aged | 92 | 19 | 83 | 100 | 30 | 324 |

## Notes

### Competing Interest Statement

The authors have declared no competing interest.

